# Matrix fibroblast function during alveolarization is dependent on GATA6

**DOI:** 10.1101/2022.06.06.494950

**Authors:** Mereena George Ushakumary, Jenna Green, Matthew Richard Riccetti, Cheng-Lun Na, Divya Mohanraj, Minzhe Guo, Anne-Karina Theresia Perl

**Author notes:** Corresponding Author, Anne-Karina Perl, PhD, Cincinnati Children’s Hospital Medical Center 3333 Burnet Avenue, MLC 7029, Cincinnati, OH 45229. Co-First Authors (Mereena George Ushakumary, Jenna Green).

## Abstract

Alveolarization is dependent on myo-, matrix- and lipo- fibroblast functions by interstitial PDGFRa^+^ fibroblasts. While these fibroblasts are derived from GLI and PDGFRa expressing fibroblasts, the transcriptional control of their functional specification remains unknown. Perinatally, the transcription factor GATA6 is upregulated in PDGFRa^+^ fibroblasts. To study the role of GATA6 during fibroblast differentiation, we generated *PDGFRa^CreER^/GATA6^flx/flx^* mice and deleted GATA6 in the perinatal period and in adult mice prior to left lobe pneumonectomy. Loss of GATA6 in the PDGFRa^+^-fibroblasts impaired alveolarization, and extracellular matrix deposition, in association with increased TCF21 expression and lipofibroblast differentiation. Loss of GATA6 in PDGFRa^+^ fibroblasts resulted in loss of alveolar type 1 (AT1) cells and gain of transitional alveolar type 2 (AT2) cells. Loss of GATA6 was associated with reduced WNT signaling. Restoration of WNT signaling in GATA6 deficient alveolar lung organoids restored AT2 and AT1 cell differentiation. GATA6 induces matrix fibroblast functions and represses lipofibroblast functions, serving as key regulator of fibroblast differentiation during alveolarization and regeneration. Present findings link matrix fibroblast functions with the ability of transitional AT2 cells to differentiate into AT1 cells.

**Graphical abstract:** Graphical abstract:

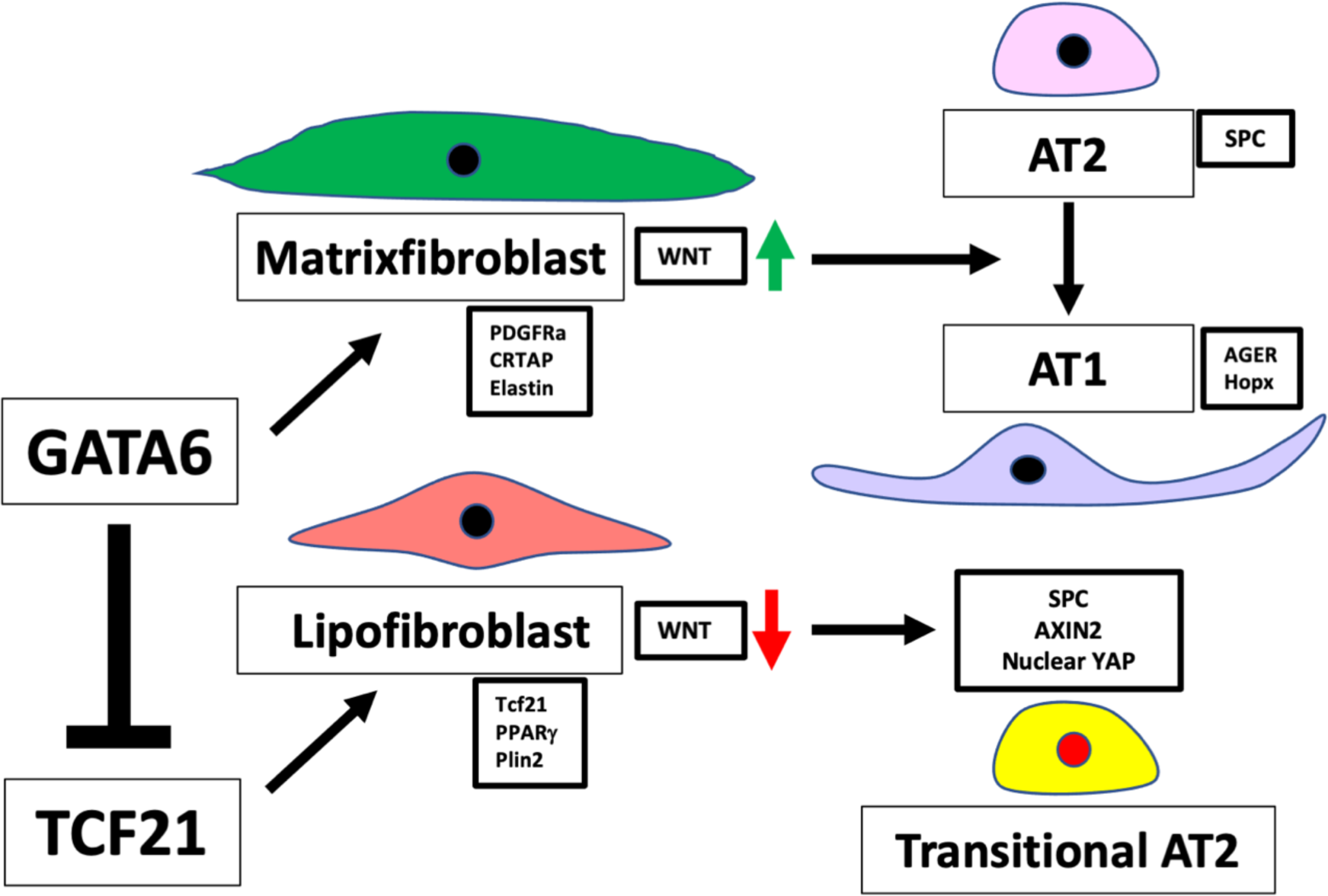

## Introduction

Alveolarization, the last stage of lung morphogenesis, occurs postnatally in mice and during the late gestational period in humans (1, 2). Premature birth and postnatal oxygen supplementation disrupt formation and thinning of secondary alveolar septa, resulting in bronchopulmonary dysplasia (BPD) (3). Lung mesenchyme has long been recognized as important in directing alveologenesis (4-9) and many mesenchymal proteins are dysregulated in human BPD and murine BPD models (10, 11). During the last decade, major advances identifying distinct fibroblast lineages, phenotypes and their roles in development and injury repair have been made (12, 13). Understanding the plasticity of lung fibroblasts and identifying key transcriptional regulators controlling their differentiation will be important in understanding pathogenic activation of fibroblasts in chronic pediatric and adult lung diseases.

Alveolar PDGFRa^+^ fibroblasts play critical roles during alveolarization and during lung regeneration (4, 14, 15). Recent studies revealed functional (myo, matrix and lipo) heterogeneities within platelet derived growth factor receptor alpha (PDGFRa^+^) fibroblasts, which change in development, disease, and regeneration (16-20). During alveolarization a subset of PDGFRA^+^ fibroblasts give rise to secondary crest myofibroblasts (SCMF) which generate the mechanical force necessary for elongation of the secondary crest as septal ridges are formed through the deposition of an elastin and collagen rich extracellular matrix (ECM) (4, 7, 21-24). Interstitial lipid laden lipofibroblasts secrete triglycerides, store retinoids, and provide trophic support for alveolar type 2 (AT2) cells (23, 25, 26). Alterations to SCMF and lipofibroblast numbers are seen in genetic and hyperoxia-induced mouse models with impaired alveolarization and in human BPD (11, 21, 22, 27). SCMF and lipofibroblasts share expression of markers like PDGFRa and glioma-associated oncogene-1 (GLI1) but are identified by unique markers e.g. alpha smooth muscle (aSMA) and Adipose Differentiation-Related Protein (ADRP). To delineate SCMF and lipofibroblast lineages multiple lineage-tracing experiments using *Pdgfra^CreERT2^, Gli1^CreERT2^, Fgf18^CreERT2^, Tcf21^mCrem^* and *Plin2^CreERT2^* have been performed (22, 28-33). SCMF are derived from lung mesenchymal progenitors during the embryonic stage that express P*DGFRa* and *GLI1* (22, 28). The number of SCMFs peaks during alveologenesis. During septal thinning SCMFs are cleared by phagocytes and are absent in the adult alveolar regions (32-34). The SCMF is characterized by the expression of aSMA which is important for secondary crest elongation during development and alveolar regeneration in adult lungs (20). SCMFs express elastin which is deposited in the alveolar septa and at alveolar entry rings to keep the alveoli open (7), little is known about other matrix organizing functions or their role in supporting AT2 cell proliferation and AT1 cell differentiation.

While PDGFRa^+^ lipofibroblasts are derived from a FGF10 lineage (28, 29, 35), their role during alveolarization is unclear. Recent lineage studies using *Gli1^CreERT^, Tcf21^mCrem^* or *Plin2^CreERT2^* demonstrated that the lipofibroblasts increase in number during alveologenesis, peaking at the end of the first phase of alveolar development and remain as a relatively stable cell population in the adult lung where they support AT2 cell function (24, 30, 31, 35, 36).

The lineage trace *Gli1^CreERT2^* marks both SCMF and lipofibroblasts suggesting a common progenitor cell (5, 28). While the SCMFs disappear at the end of alveolarization, lipofibroblasts persist in adult murine lungs. Injury in the adult lung can induce a lipo to myo fibroblast transition, suggesting a functional switch or injury dependent fibroblast activation (37). Moreover, attenuation of lipo-to-myo-fibroblast differentiation during injury reduced ECM deposition, suggesting a presently poorly understood matrix remodeling function, which is independent of the contractile function of myofibroblasts (27, 38-40). Single cell RNA sequencing (ScRNA seq) studies have identified an alveolar matrix fibroblast population defined by the expression of matrix associated genes and lack of contractile function. These cells express PDGFRa^+^, but their lineage relationship to myo- and lipofibroblasts remains unclear (41). Recent studies demonstrate that loss of matrix fibroblast function with advanced age and in idiopathic pulmonary fibrosis results in impaired alveolar regeneration and impaired AT1 differentiation and is associated with fragmented ECM (42). In a perinatal hyperoxia model the number of PDGFRa^+^ fibroblasts was reduced and PDGFRa^+^ fibroblasts switched from a myo to a lipo phenotype in association with reduction in matrix organizing function and lack of AT1 differentiation (43). The molecular regulator controlling matrix fibroblasts and their specific role in alveolarization are unknown.

Integration of developmental ScRNA seq datasets from LungMAP, bulk-RNA seq from PDGFRa^+^ FB populations isolated from lungs during alveolar septation, and bulk-RNA seq from lungs in a hyperoxia model of impaired alveolar regeneration predicted GATA6 as a key regulator of myo and matrix- specification in PDGFRa^+^ fibroblasts (16-18, 42). While GATA6 expression is lost in aged fibroblasts associated with loss of matrix function (42), expression of GATA6 is increased in PDGFRa^+^ fibroblasts during peak alveolarization (44). To specifically inactivate GATA6 in emerging SCMF we used *PDGFRa^CreERT2^* and GATA6^flx/flx^ mice and inactivated GATA6 by administrating a single intraperitoneal tamoxifen dose to PN1 pups. Loss of GATA6 in PDGFRa fibroblasts (GATA6^PDGFRAΔ/Δ^) caused alveolar simplification, loss of extracellular matrix and increased lipofibroblast differentiation. Loss of GATA6 in SCMFs resulted in an increase in AT2 cells with increased Axin2 and nuclear YAP suggesting a halt of AT2 to AT1 differentiation (45, 46). In summary these data suggest an important role of SCMF in ECM remodeling and support of alveolar epithelial differentiation in the alveolar niche.

## Results

### Postnatal inactivation of GATA6 in SCMF impairs alveolarization and extracellular matrix deposition

We generated PDGFRa^CreER^/GATA6^flx/flx^ mice and deleted GATA6^PDGFRAΔ/Δ^ using one dose of tamoxifen on PN1. Lungs were obtained on PN7 and PN28 for gene expression analysis, histology and transmission electron microscopy (Fig. 1A). GATA6 mRNA in isolated PDGFRa^+^ fibroblasts was significantly decreased at PN7 but did not reach statistical significance at PN28, suggesting that GATA6 was successfully inactivated in SCMF which by PN28 have been cleared (Fig. 1B). Histology of multiple biological replicates at PN7 and PN28, revealed alveolar simplification (Fig. 1C). Quantitative morphometry using Mean Linear Intercept (Lm), Volume Density of alveolar septa (Vvsep), Mean Transsectional wall length (LMW) and Surface area density of airspace (SVair) showed significantly reduced Vvsep, Lmw at both timepoints and increased SVair at PN7 (Fig. 1D). Weigert’s elastin staining indicated less elastin (black) and collagen (pink/red) deposition in the septal walls of GATA6^PDGFRAΔ/Δ^ lungs at PN7 (Supplemental Figure 1A). Since our previous work demonstrated an important role of PDGFRa+ fibroblasts in realveolarization after partial pneumonectomy (PNX) (20), we tested the role of GATA6 in this lung regeneration model. Control and experimental mice were treated with tamoxifen three days prior to PNX to delete GATA6. Lungs were harvested at Day 5 post-PNX. Histological studies demonstrate alveolar simplification, suggesting impaired regeneration (Supplement Fig. 1B).

**Figure 1:**
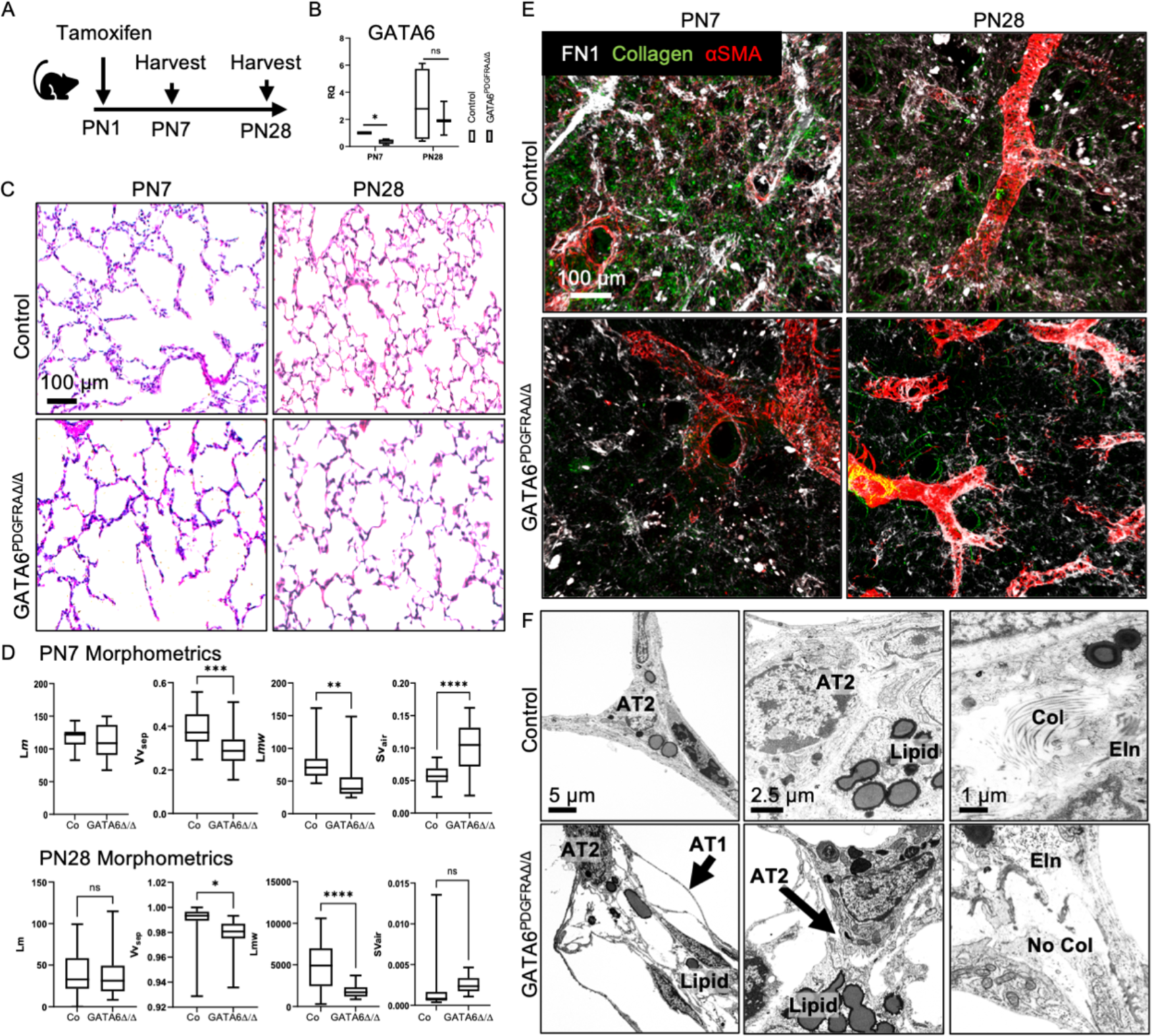
Deletion of GATA6 in PDGFRA+ fibroblasts results in alveolar simplification and loss of extracellular matrix. **(A)** Timeline and tamoxifen treatment of mice used in the study. **(B)** MACS-isolated PDGFRA^+^ fibroblasts show reduced GATA6 gene expression by RT-qPCR, *n* = 3-6 Control and GATA6^PDGFRΔ/Δ^ mice. Data displayed as box-and-whisker plot. The box plots depict the minimum and maximum values (whiskers), the upper and lower quartiles, and the median. One-way ANOVA was used, *p<0.05. Error bars show mean ± SD. **(C)** H&E staining of Control and GATA6^PDGFRΔ/Δ^ mice at PN7 and PN28. Scale bar = 100 μm. **(D)** Alveolar Morphometrics at PN7 & PN28, Graphs represents Lm (mean linear intercept of airspaces), Vvsep (volume density of alveolar septa), Lmw (Mean transsectional wall length) and Svair (Surface area density of airspaces). Two-tailed Student’s t-test was used, ***p<0.0005, **p<0.005, ****p<0.0001, *p<0.05. **(E)** Second harmonic imaging for collagen and immunohistochemistry for fibronectin and alpha smooth muscle at PN7. Scale bar = 100 μm. **(F)** Transmission electron microscopy of Control & GATA6^PDGFRAΔ/Δ^ lungs at PN7 showing detachment of AT1 and AT2 cells from ECM, increased lipid droplets (Lipid) and reduced collagen (Col), Arrows pointing at detached epithelial cells. Scale bar = 5μm, = 2.5 μm, =1μm.

Confocal imaging using second harmonic generation for collagen and immunohistochemistry for fibronectin and alpha smooth muscle demonstrated reduction of fibronectin and loss of fibrillar collagen in GATA6^PDGFRAΔ/Δ^ lungs at PN7 (Fig. 1E). Transmission electron microscopy of GATA6^PDGFRAΔ/Δ^ lungs at PN7 demonstrated detachment of AT1 and AT2 cells from the ECM, increased lipid droplets in fibroblasts and fragmented collagen bundles (Fig. 1F). Collagen fibers were restored and AT1 cells reattached to basal membranes by PN28 (Supplemental Fig. 1C). The decrease in collagen can be attributed to the increase in Cathepsin K gene expression in the isolated PDGFRa+ cells (Supplemental Fig. 1D ). These data imply a critical regulatory role of GATA6 for matrix function in SCMF.

While the SCMF generates the contractile force to elongate the newly forming septa additional ECM needs to be synthesized to provide structural support. Cartilage-associated protein (CRTAP) and Fibronectin (FN1), both proteins made by matrix fibroblasts were reduced in GATA6^PDGFRAΔ/Δ^ lungs at PN7 (Fig. 2 A, B), but the RNA expressions of these proteins were unchanged on PN7 (Supplemental Fig. 2B). Similarly CRTAP and FN1 were reduced on day 5 post-PNX (Supplemental Fig. 2A). While collagen bundles and FN1 were restored by PN28, CRTAP remained reduced in PN28 lungs (Supplemental Fig. 2C, D). Taken together GATA6 plays an important role in SCMF matrix fibroblast function during alveolarization and re-alveolarization after PNX.

**Figure 2:**
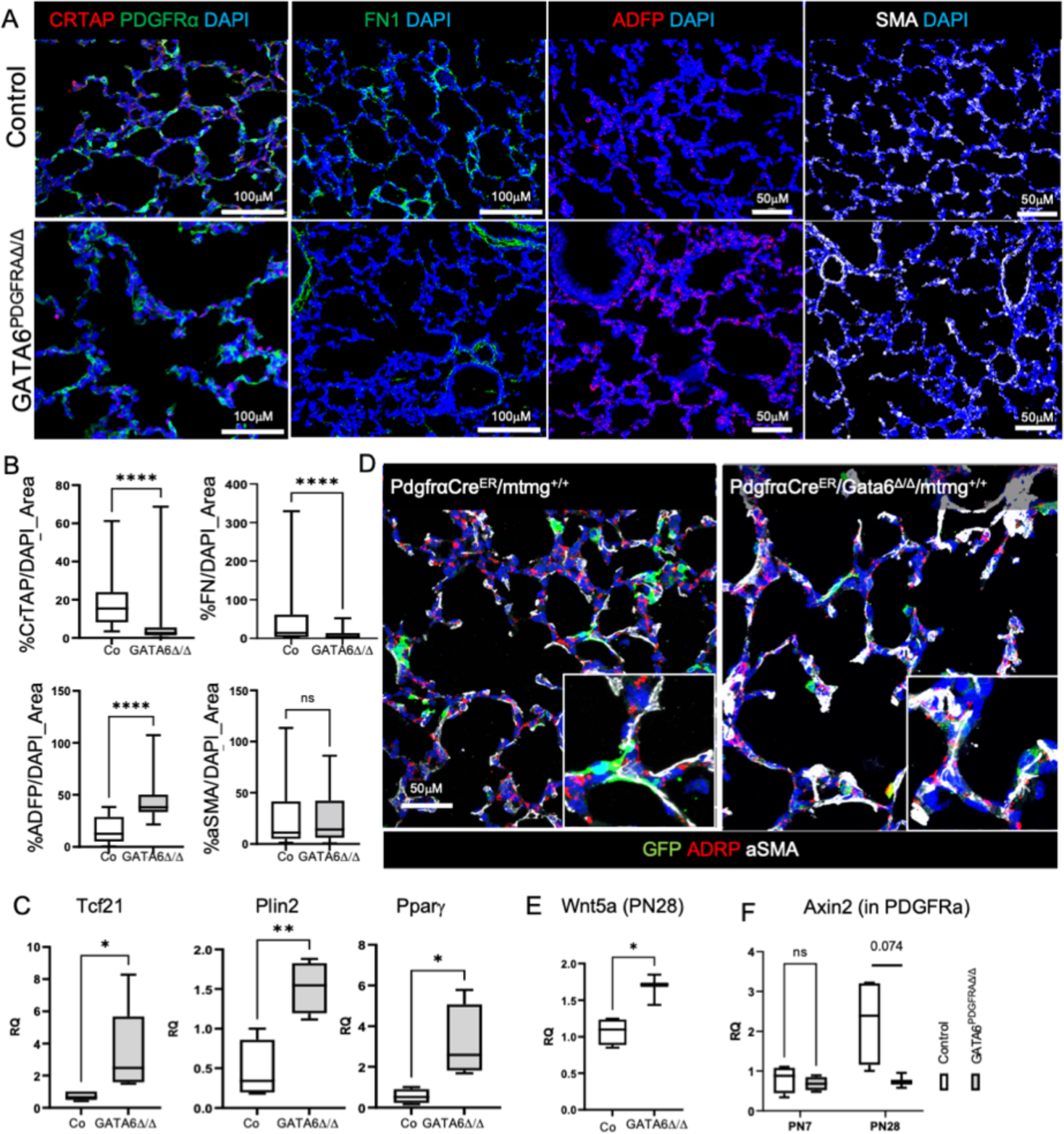
GATA6 inhibits lipofibroblast differentiation and regulates matrixfibroblast function. **(A)** PN7 Control and GATA6^PDGFRAΔ/Δ^ immunofluorescence for matrix (CRTAP, FN1), lipo (ADRP) and myo (αSMA) fibroblast markers. CRTAP and FN1 were reduced in GATA6^PDGFRAΔ/Δ^ lungs, ADRP was increased and αSMA was unchanged **(B)** Quantification of CRTAP, FN1, ADRP and αSMA with Nikon Elements as antibody area over DAPI area, n=3-6. Two-tailed Student’s t-test, ****p<0.0001, ns-not significant **(C)** q- RTPCR of MACS-sorted PDGFRa+ fibroblasts: Increased expression of Tcf21, Plin2 and Pparγ in GATA6^PDGFRAΔ/Δ^ fibroblasts at PN7. n=3-6, Two-tailed Student’s t-test, *p<0.05, **p= 0.0053. **(D)** Immunofluorescence for PDGFRa-GFP lineage trace of SCMF and colocalization of aSMA (white) and ADRP(red) in PN7 control and GATA6^PDGFRAΔ/Δ^ lungs. Efficient recombination in control and GATA6^PDGFRAΔ/Δ^ lungs targets myo and lipofibroblasts. **(E)** Increase in Wnt5a expression in MACS-sorted PDGFRa+ fibroblasts of GATA6^PDGFRAΔ/Δ^ lungs at PN28. n=3-4, Two-tailed Student’s t-test, *P<0.05. **(E)** Increase in Axin2 gene expression at PN28, n=3-4, One-way ANOVA was used. p=0.074 at PN28.

### Inactivation of GATA6 in SCMF increases numbers of lipofibroblasts but does not change myofibroblast differentiation

Perinatal hyperoxia exposure decreased matrix and increased lipo fibroblast function in PDGFRA^+^ fibroblasts (43). Lipofibroblast differentiation was assessed by immunofluorescence staining for adipose differentiation-related protein (ADFP). Lipofibroblast numbers were significantly increased at PN7 (Fig. 2 A, B) and 5 days after PNX (Supplemental Fig. 2 A). The increase in lipofibroblasts was not sustained through PN28 (Supplement Fig. 2 C, D). Expression of lipofibroblast signature genes, Tcf21, Plin2 and Pparψ, in isolated PDGFRa^+^ fibroblasts was significantly increased at PN7 (Fig. 2C). These data suggest that GATA6 suppresses lipofibroblast differentiation within the PDGFRA+ population during alveolarization and regeneration.

Lineage tracing using a mT/mG reporter demonstrate efficient targeting of myo and lipofibroblasts in control and GATA6^PDGFRAΔ/Δ^ lungs at PN7 (Fig.2 D). At PN28 lineage traced GATA6^PDGFRAΔ/Δ^ are cleared by macrophages and lineage traced fibroblasts do not co-stain with aSMA or ADRP (Fig.2 E). Immunofluorescence staining for αSMA revealed that myofibroblast differentiation was unchanged at PN7 (Fig. 2A, B). Increased myofibroblast differentiation was detected at PN28 (Supplement Fig. 2 C, D), suggesting activation of myofibroblast function in non SCMFs. Myofibroblast differentiation in non SMCFs was accompanied with an increase in Wnt5a gene expression and a decrease in Axin2 (Fig. 2 F).

### GATA6 suppresses lipofibroblast function in human lung fibroblasts in vitro

Immortalized human fetal lung fibroblasts (IMR90) express PDGFRa and GATA6. To determine if GATA6 is directly regulating lipo, matrix and myofibroblast gene expression, IMR90 cells were stably transfected with lentiviral GATA6 shRNA and GATA6 expression plasmids. GATA6 shRNA transfection efficiency and GATA6 overexpression was confirmed by qPCR and western blot (Supplemental Fig. 3A, B). Inactivation of GATA6 in IMR90s significantly increased the lipofibroblast associated transcription factor TCF21 and PLIN2 and significantly decreased matrix fibroblast signature genes ELN, expression of ACTA2 and COL1A1 did not change significantly (Fig. 3A). WNT2A was significantly inhibited by GATA6 shRNA and induced by expression of GATA6 (Fig. 3A), suggesting that GATA6 directly regulates WNT2. Inhibition of GATA6 decreased contraction of collagen pellets by 25.24% (Fig. 3B) (control mean = 0.462 cm2, shRNA mean = 0.618 cm2, p < 0.001, GATA6 OE mean = 0.489 cm2, p > 0.05). Immunofluorescence for F-actin and second harmonic imaging of the collagen pellet showed decreased collagen after inhibition of GATA6 and increased F-Actin stress fibers after GATA6 induction (Fig. 3C). These data demonstrate that GATA6 suppresses lipo function and induces matrix function while the myo function remains largely unchanged.

**Figure 3:**
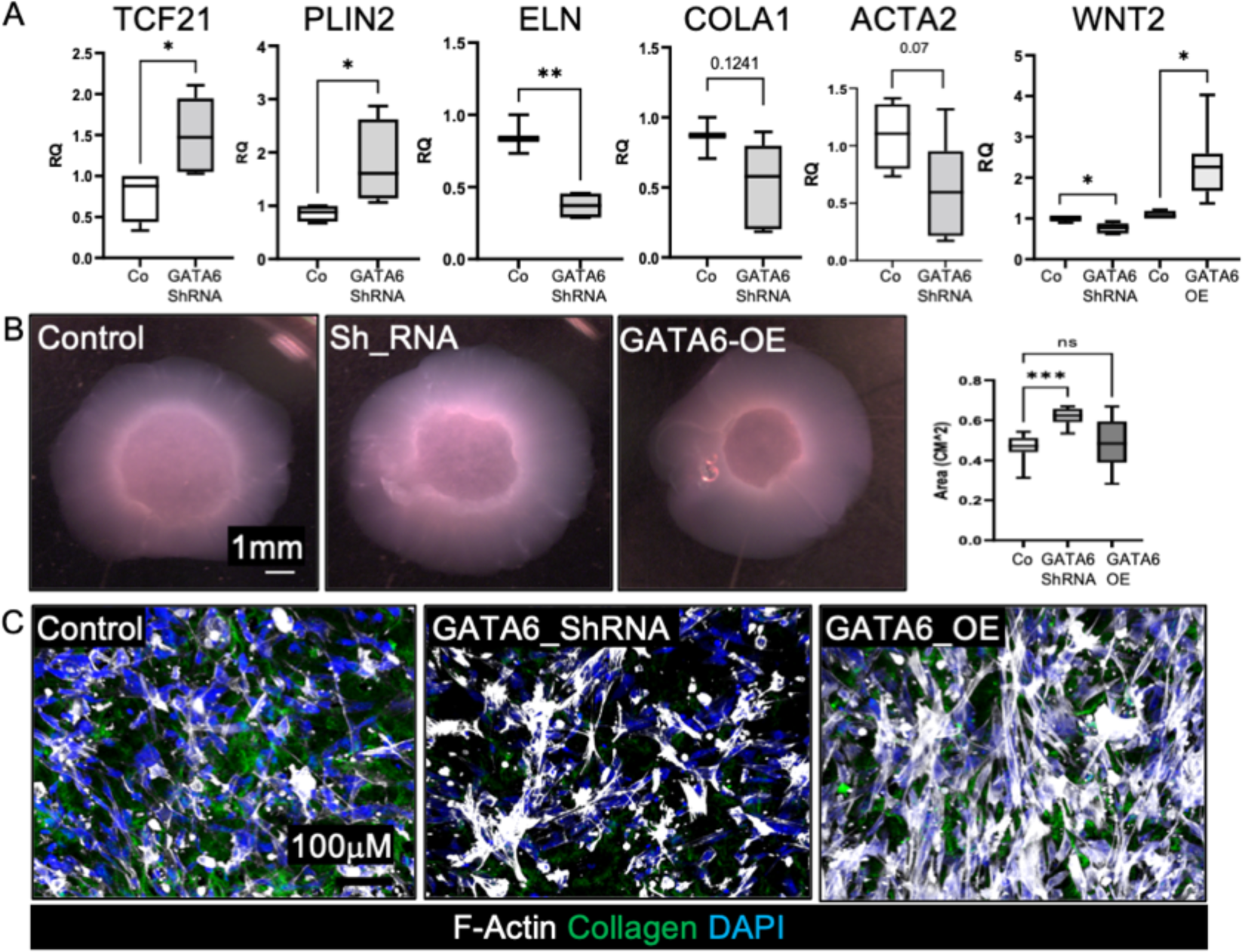
GATA6 suppresses lipofibroblast and increases matrixfibroblast differentiation in human lung fibroblasts. **(A)** Stable transfection of IMR90 cells with GATA6 ShRNA significantly increased TCF21 and PLIN2 expression, and significantly reduced matrix fibroblast marker ELN. Expression of COLA1 and ACTA2 did not change significantly. n=3-6, Two-tailed Student’s t-test, *p<0.05, **p<0.005, ns- not significant. GATA6 ShRNA significantly reduced WNT2 expression and GATA6. Overexpression increased WNT2 gene expression n=3-8, One-way ANOVA was done. *p<0.05. (**B)** Collagen contraction assay with GATA6 ShRNA, GATA6-OE and scrambled ShRNA Control IMR90s after 24 hrs in culture. Quantification of contraction was measured as area in CM^2. GATA6 ShRNA-treated IMR90s contracted collagen less compared to controls. n = 12, Ordinary One-way ANOVA, ***P < 0.0005, ns-not significant. **(C)** F-actin immunofluorescence and second harmonic generation of collagen pellets. Decreased collagen in GATA6 ShRNA IMR90s and increased in f-actin stress fibers in GATA6 OE IMR90s. Scale bar=100µM.

### GATA6, EZH2 and TCF21 work in concert to regulate lipofibroblast differentiation

TCF21 is known to regulate lipofibroblast differentiation in the lung (31). TCF21 mRNA was increased in primary GATA6^PDGFRAΔ/Δ^ fibroblasts and GATA6 shRNA transfected IMR90 fibroblasts suggesting that TCF21 was regulated by GATA6 (Fig. 2C and Fig. 3A). In small cell lung cancer and clear cell sarcoma of the kidney TCF21 is inhibited by EZH2 mediated hypermethylation (47, 48). EZH2 is a transcriptional repressor which is part of the polycomb repressive complex2 (PCR2). EZH2 like GATA6, is increased in PDGFRa^+^ fibroblasts during alveolarization, suggesting a role for EZH2 in suppressing lipofibroblast differentiation (Supplemental Fig. 3C-D). ShRNA-mediated inhibition of GATA6 decreased EZH2 protein (Fig. 4A). Inhibition of EZH2 with the methyltransferase inhibitor 3-Deazaneplanocin (DZNep) in IMR90 increased TCF21 expression but did not change GATA6 expression (Fig. 4B). These data suggest that GATA6 regulates lipofibroblast specification through a mechanism involving EZH2 and TCF21. Chromatin immunoprecipitation using EZH2 antibody and quantitative RT-PCR of predicted binding sites in the TCF21 promoter were performed (ChIP-PCR). EZH2 bound to the TCF21 promoter and was significantly increased by overexpression of GATA6 in IMR90s. EZH2 binding to the TCF21 promoter in GATA6-shRNA transfected IMR90s was not significantly altered (Fig. 4C). These data suggest that GATA6 inhibits TCF21 via increased EZH2 binding to the TCF21 promoter. Inhibition of TCF21 via shRNA increased GATA6 gene expression suggesting a positive feed back loop of TCF21 on its own expression via inhibition of GATA6 (Fig. 4D). In summary these data suggest that GATA6 and EZH2 suppress lipofibroblast differentiation via transcriptional control of TCF21 (Fig. 4E).

**Figure 4:**
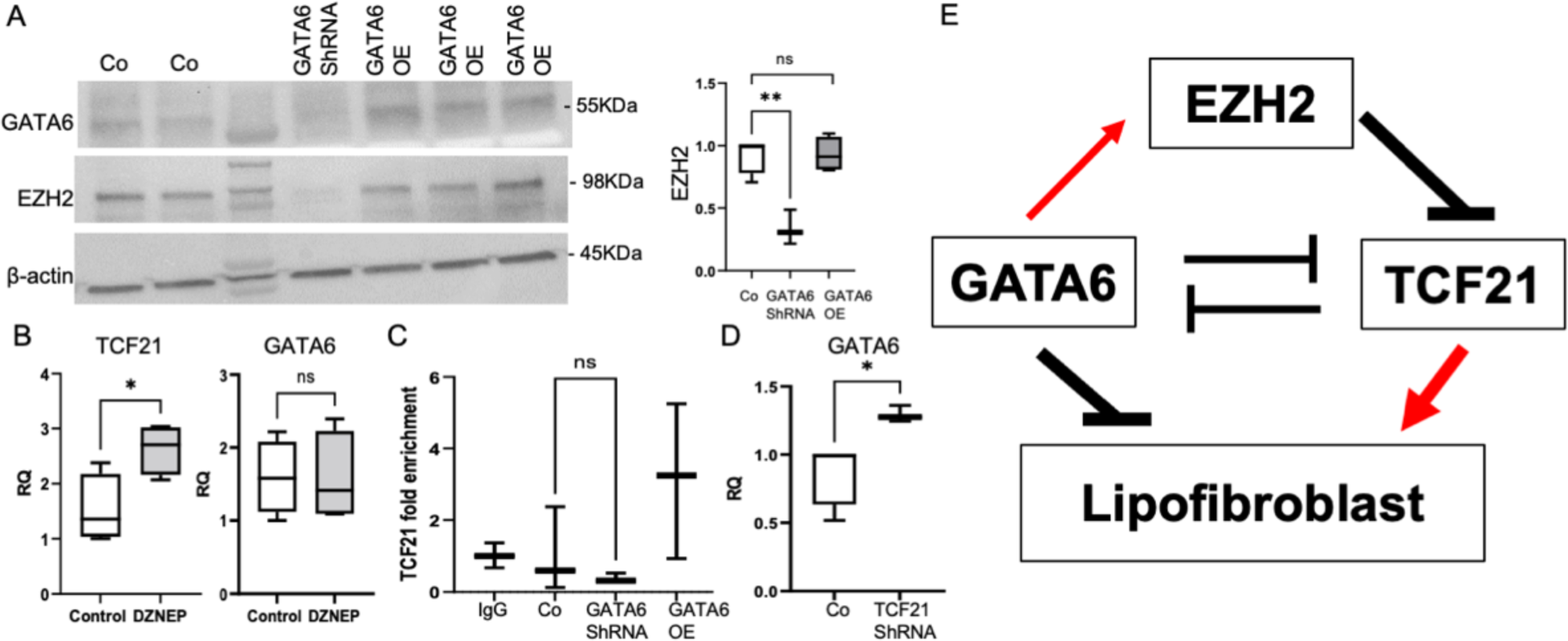
GATA6 and EZH2 suppress TCF21 expression. **(A)** Western protein blot of stable transfected GATA6 ShRNA and GATA6_OE IMR90s: Significant reduction of GATA6 and EZH2 protein in GATA6 shRNA IMR90s. β-actin: loading control. EZH protein quantification by densitometric analysis of scanned blots using ImageJ software. One-way ANOVA. n=3-5, **p<0.005, ns-not significant. **(B)** Inhibition of EZH2 with DZNEP in IMR90s increased TCF21 but not GATA6 gene expression. n=3-4, Two-tailed Student’s t-test was used, *P<0.05, ns-not significant. **(C)** Chromatin Immunoprecipitation using EZH2 antibodies and primers for TCF21 promoter. EZH2 binding to the TCF21 promoter was significantly increased in GATA6 Over expression plasmid-transfected IMR90s. n=3, One-way ANOVA. ns-not significant. **(D)** ShRNA knock down of TCF21 increased GATA6 gene expression in IMR90s. n=3-4, Two- tailed Student’s t-test, *P<0.05. **(E)** Schematic regulation of lipofibroblast differentiation by GATA6 suppressing TCF21 via EZH2.

### Loss of matrix fibroblast function results in AT2 hyperplasia and increase in transitional AT2/AT1 cells

PDGFRA^+^ fibroblasts support alveolar type 2 (AT2) and alveolar type 1 (AT1) cell differentiation during alveolarization, homeostasis and injury repair (15, 42, 43, 49, 50). Immunohistochemistry for the AT2 cell marker proSPC and AT1 cell markers HOPX and AGER were performed on control and GATA6 ^PDGFRAΔ/Δ^ lungs at PN7 and PN28 (Fig. 5A, C). Numbers of SPC positive cells were increased and HOPX and AGER positive cells decreased in GATA6 ^PDGFRAΔ/Δ^ lungs at PN7 (Fig. 5B). Similarly, at PN28 numbers of SPC positive AT2 were increased and HOPX positive AT1 were decreased. Numbers of AGER positive AT1 were not significantly decreased at PN28 (Fig. 5D). Likewise, numbers of SPC positive AT2 cells were increased and AGER positive AT1 cells decreased during lung regeneration after PNX (Supplemental Fig. 4A). These data suggest that GATA6 regulates SCMF functions which are important for AT2 to AT1 differentiation. During normal lung development NKX2.1 and YAP regulate AT1 specification in transitional AT1/AT2 cells (51). After deletion of GATA6 in fibroblasts nuclear YAP was increased in SPC positive AT2s at PN28 (Fig. 5E), but not at PN7 (Supplemental Fig. 4B). Axin2 positive AT2 progenitor cells are important for alveolar regeneration (45). Axin2 mRNA was increased in alveolar epithelial cells isolated from PN7 and PN28 GATA6^PDGFRAΔ/Δ^ lungs (Fig. 5F). These data demonstrated that in GATA6 ^PDGFRAΔ/Δ^ lungs AT1 differentiation is impaired resulting in increased numbers of transitional AT2/AT1 or progenitor AT2 cells. These data indicate that GATA6 regulates the ability of SCMFs to support AT1 specification.

**Figure 5:**
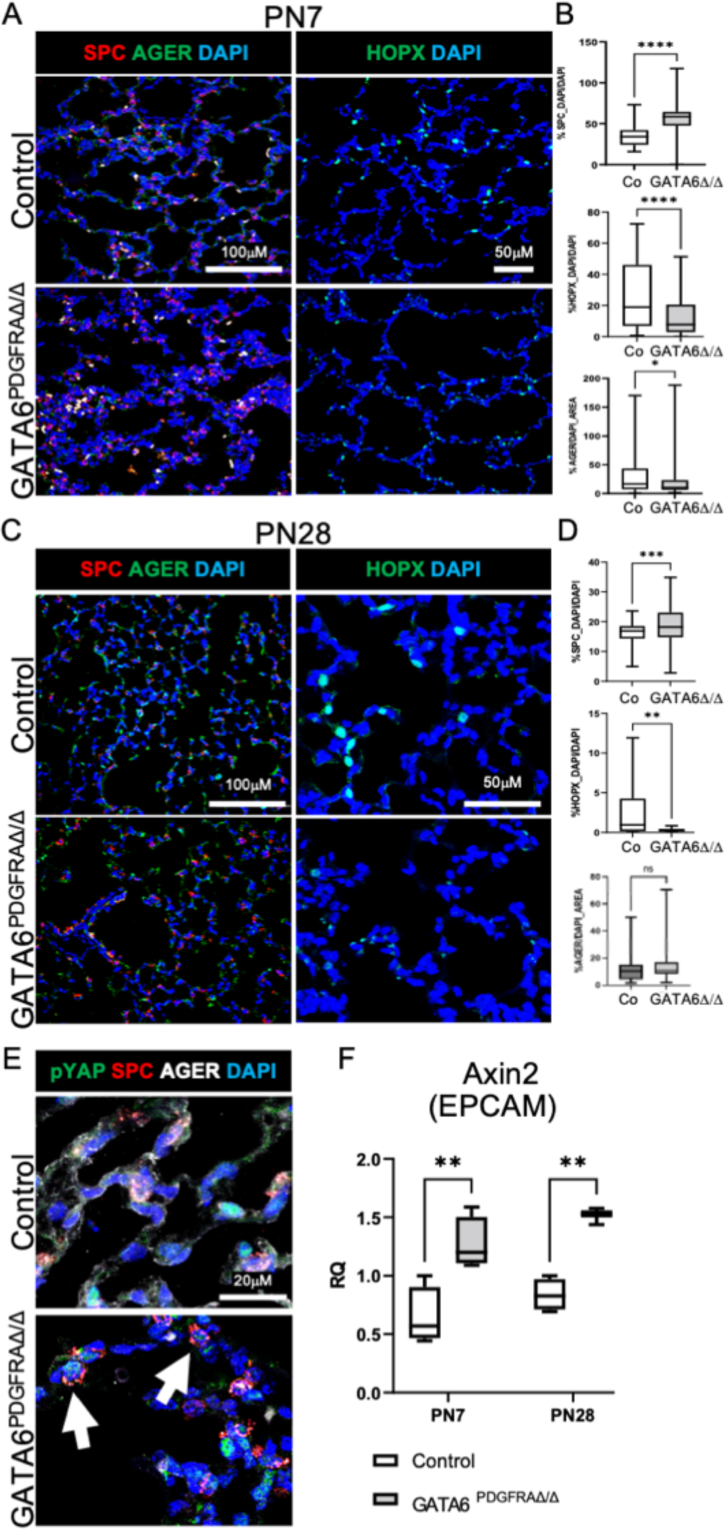
Deletion of GATA6 in SCMFs impairs AT2 to AT1 differentiation. **(A)** PN7 Control and GATA6^PDGFRAΔ/Δ^ immunofluorescence for AT1 (HOPX, AGER) and AT2 (proSPC) cell markers: Increase in SPC and decrease in HOPX and AGER positive cells in GATA6^PDGFRAΔ/Δ^ lungs. scale bars= 100 µM & 50 µM **(B)** Quantification of PN7 immunofluorescence with Nikon Elements show significant increase in SPC and decrease in HOPX and AGER. Nuclear antibody stain HOPX and SPC fluorescence were quantified over DAPI, and cytoplasmic and membrane stains (AGER,) as % area over DAPI area. n=3-6, Two-tailed Student’s t-test was used, ****p<0.0001, *p<0.05. **(C)** PN28 Control and GATA6^PDGFRAΔ/Δ^ immunofluorescence for AT1 (HOPX, AGER) and AT2 (proSPC) cell markers: Increase in SPC and decrease in HOPX positive cells in GATA6^PDGFRAΔ/Δ^ lungs., scale bars= 100 µM & 50 µM **(D)** Quantification of PN28 immunofluorescence with Nikon Elements show significant increase in SPC and decrease in HOPX. Nuclear antibody stain HOPX and SPC fluorescence were quantified over DAPI, and cytoplasmic and membrane stains (AGER,) as % area over DAPI area. n=3-6, Two-tailed Student’s t-test was used, ***p<0.0005, **p<0.005, ns= not significant. **(E)** Colocalization of YAP and SPC immunofluorescence was detected in GATA6^PDGFRAΔ/Δ^ lungs at PN28 (Arrow). Nuclear YAP in control lungs did not colocalize with SPC. scale bars= 20 µM **(F)** Axin2 mRNA expression in MACS-Sorted epithelial cells isolated from GATA6^PDGFRAΔ/Δ^ lungs was significantly increased at PN7 and PN28. n=3-4, 2-way ANOVA. **p< 0.005.

### GATA6^PDGFRAΔ/Δ^ fibroblasts fail to support alveolar epithelial differentiation in vitro

Primary PDGFRA^+^ fibroblasts support in vitro lung organoid formation and can be used to interrogate epithelial mesenchymal interactions that promote organoid formation, growth and AT2/AT1 differentiation (42, 43, 52). PDGFRA^+^ cells from PN7 control and GATA6^PDGFRAτι/τι^ lungs were cultured with normal CD326 sorted epithelial cells from adult mouse lungs. After 3 weeks in organoid cultures expression for E-Cadherin, SPC, AGER, HOPX, PDGFRa and aSMA was assessed by immunohistochemistry. Control fibroblasts formed small alveospheres with SPC positive cells on the outside and AGER positive cells on the inside (Fig. 6A). GATA6^PDGFRAτι/τι^ fibroblasts formed larger spheres with few positive SPC cells, expression of AGER and HOPX was not detected (Fig. 6B). In control organoids epithelial cells were surrounded by aSMA positive fibroblasts, but in the GATA6^PDGFRAτι/τι^ organoids, epithelial cells were not in contact with aSMA positive fibroblasts. A thin layer of PDGFRa^+^ fibroblasts was associated with epithelial cells in control and GATA6^PDGFRAτι/τι^ organoids. Taken together these data demonstrate that GATA6^PDGFRAτι/τι^ PDGFRA^+^ fibroblasts fail to support epithelial differentiation in organoids.

**Figure 6:**
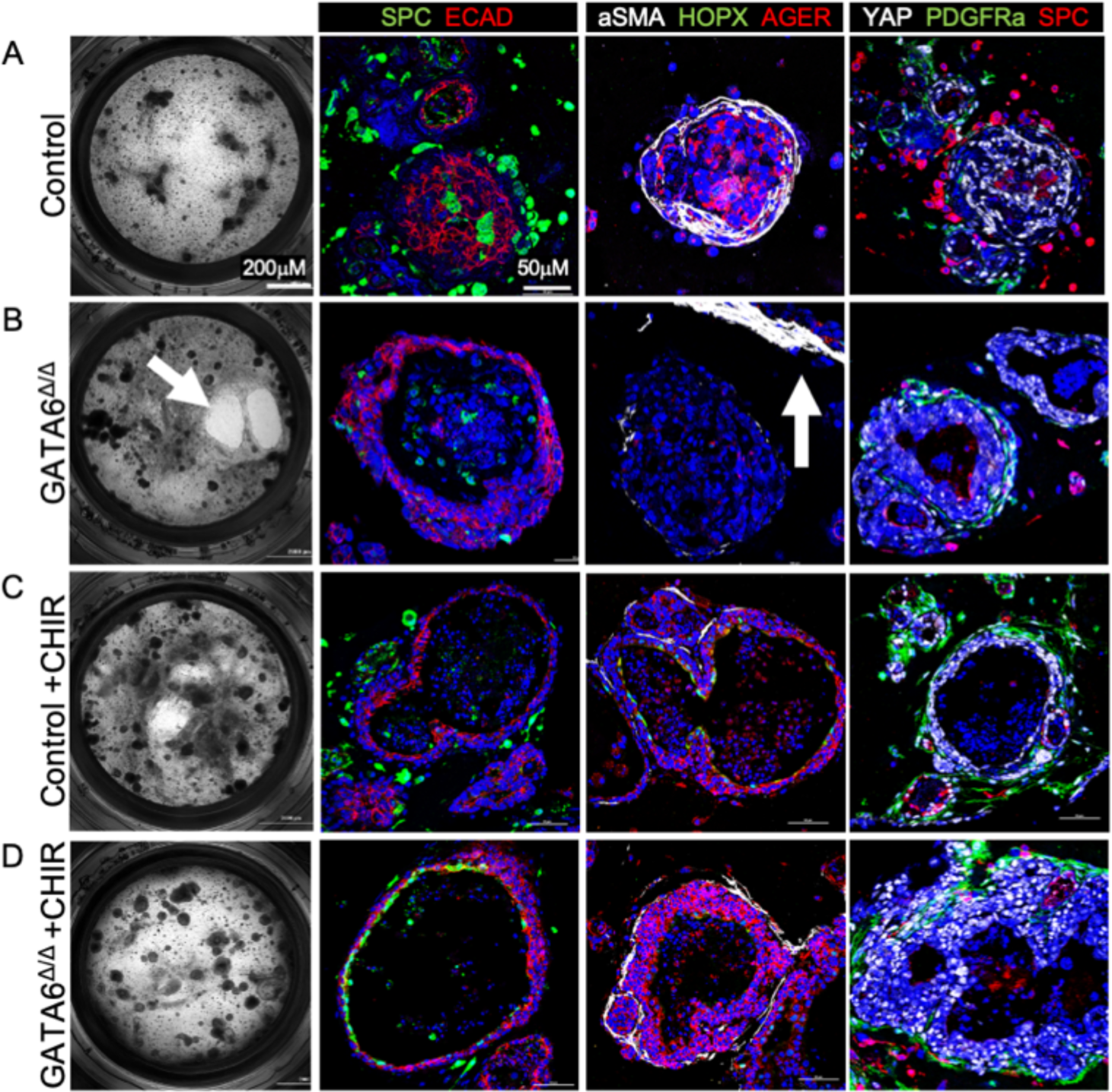
Gata6 deficient fibroblasts do not support epithelial differentiation in organoids which is restored by CHIR. Fibroblasts from PN7 control and GATA6^PDGFRAΔ/Δ^ lungs were cocultured with adult lung epithelial cells for 3 weeks and imaged by Bright-field. Scale bar =200µM. N=3-10 organoid transwells (replicates), 3 slides per transwell. Immunofluorescence on paraffin sections for epithelial and fibroblast markers epithelial differentiation. Scale bar =50µM. **(A)** Organoids with control fibroblasts formed small alveolospheres with SPC positive cells on the outside and AGER positive cells on the inside. aSMA positive fibroblasts and a thin layer of PDGFRa fibroblasts surround the spheres. Nuclear Yap is found in non SPC epithelial cells. **(B)** Cultures with GATA6^PDGFRAΔ/Δ^ fibroblasts degraded the Matrigel and formed large organoids with few SPC cells inside, AGER and HOPX was not detected. aSMA fibroblasts were not in contact with epithelial cells (Arrow). **(C)** CHIR treatment of control organoids did not change epithelial differentiation **(D)** CHIR treatment of organoids with GATA6^PDGFRAΔ/Δ^ fibroblasts restored SPC and AGER expression in epithelial cells and contact of aSMA positive fibroblasts with epithelial cells. CHIR reduced Matrigel increased nuclear YAP in epithelial cells and increased the layer of PDGFRA fibroblasts organoids.

### Deletion of GATA6 in fibroblasts impairs WNT signaling in both fibroblasts and epithelial cells

Epithelial expression of GATA6 has been extensively studied and epithelial specific GATA6 deletion in lung epithelial cells and cardiomycytes identified a role of GATA6 in regulating Wnt signaling (53-58). To determine dynamic changes in gene expression within the PDGFRA^+^ population and alveolar epithelial cells, we performed bulk RNA-seq on sorted PDGFRA^+^ cells and EPCAM^+^ epithelial cells from control and GATA6^PDGFRAτι/τι^ PN7 murine lungs. Gene ontology (GO) analysis on differentially expressed genes (FC>2 and p value<.01) was performed and confirmed activation of a pathological fibroblast and intermediate AT2 cells, similar to AT2 cells found in idiopathic pulmonary fibrosis (Supplemental Fig. 5A-C). A heatmap for WNT signaling in PDGFRa^+^ and epithelial cells was generated with 152 KEGG WNT pathway genes. The majority of genes were differentially expressed between fibroblasts and epithelial cells. In the fibroblasts 36 WNT associated genes were differentially expressed suggesting downregulation of canonical WNT signaling (Supplemental Fig. 5D). WNT pathway genes were validated by qPCR in primary fibroblasts, IMR90s and epithelial cells (Fig. 2, 3, 5). To assess whether activation of canonical Wnt signaling restores AT2/AT1 differentiation in GATA6^PDGFRAτι/τι^ organoids, we cultured organoids in the presence of CHIR (Fig. 6C, D). CHIR treatment of control organoids did not change epithelial differentiation (Fig. 6C). CHIR treatment of organoids made with GATA6^PDGFRAτι/τι^ fibroblasts restored SPC expression in epithelial cells lining the organoid lumen and AGER expression in cells on the external surface of the organoids (Fig. 6D). CHIR arestored the contact of aSMA^+^ fibroblasts to epithelial spheres and reduced Matrigel degradation. CHIR increased nuclear YAP in epithelial cells of GATA6^PDGFRA,1/,1^ organoids and increased the layer of PDGFRa^+^ fibroblasts around control and GATA6^PDGFRA,1/,1^ organoids. These data demonstrate that activation of WNT signaling via CHIR restored the ability of GATA6^PDGFRA,1/,1^ fibroblasts to support AT2 and AT1 differentiation.

## Discussion

We used loss-of-function genetics, in vitro gain and loss of function, and 3D organoid cultures to elucidate the role of GATA6 in regulating matrix and lipofibroblast function in PDGFRa+ SCMFs. Inactivation of GATA6 in SCMFs caused loss of extracellular matrix and gain of lipofibroblast function and impaired alveolarization and AT1 differentiation. ChIP-PCR and methyltransferase inhibition support the concept that GATA6 and EZH2 bind and regulate the TCF21 promoter, suppressing lipofibroblast differentiation. Change in fibroblast differentiation resulted in an increase of Axin2 positive AT2 cells expressing nuclear YAP, previously referred to as “alveolar progenitors” or “transitional AT2 cells”. These findings support a model in which GATA6 regulates matrix fibroblast differentiation which in turn regulating AT2 to AT1 epithelial transition and alveolarization. Using GATA6 deficient fibroblasts in organoid cultures, we show that activation of WNT signaling restores the capacity of GATA6 deficient fibroblasts to support AT1 differentiation.

### GATA6 regulates cell differentiation and function

GATA6 is a zinc finger transcription factor that works in concert with other transcription factors to regulate cell differentiation and is expressed in endo- and mesoderm derived cells (59-61). Epithelial expression of GATA6 has been extensively studied and epithelial specific GATA6 deletion increased Wnt signaling which impaired distal epithelial differentiation and alveolar septation (53-57). The process of cardiogenesis is regulated by GATA transcription factors in tandem with Wnt signaling (58). Deletion of GATA6 in cardiomyocytes resulted in fundamental changes in the levels of key regulatory genes and myocyte differentiation-specific genes while increased expression caused pathological cardiac hypertrophy (61-64). Here, we show that GATA6 expression in PDGFRa fibroblasts is required to suppress lipofibroblast differentiation and to regulate paracrine signaling supporting proper alveolar epithelial differentiation in mice. Present GATA6 gain and loss of studies in IMR90s show a direct correlation of WNT2a and GATA6. Activation of WNT by treating GATA6^PDGFRA,1/,1^ fibroblast organoids with CHIR restored their ability to support AT1 differentiation. Together these data demonstrate that GATA6 regulates matrix and lipofibroblast differentiation and WNT signaling required for AT2 to AT1 epithelial cell differentiation during alveolarization.

### Novel matrix function of the SCMF

Lung development and regenerative responses are regulated by functionally and spatially by distinct fibroblast subpopulations. PDGFRa+ alveolar fibroblasts are vitally important for alveolar septation and epithelial cell differentiation (17, 65, 66). This group of fibroblasts comprises three functional stages (myo, lipo and matrix) whether these fibroblasts change functional stages or are independent linages/populations needs to still be determined. The present study we focused on the transcriptional regulation of matrix and lipofibroblast function within the SCMFs. Perinatal deletion of GATA6 from PDGFRa+ fibroblasts did not change the contractile function and expression of aSMA, but significantly increased lipofibroblasts differentatition based on increased expression of ADFP, Plin2, Tcf21 and Pparψ. In addition to the loss of matrix proteins e.g. Crtap and fibronectin we identified a novel collagenolytic function of lipofibroblasts mediated by upregulation of cathepsins. Electron microscopy and 3D confocal reconstruction demonstrated loss of ECM and degradation of fibrillar collagen. The collagenolytic function of the fibroblasts was inhibited by GATA6, providing a new strategy for resolution of fibrosis for future investigations using loss-of function models and integration of gene expression studies and proteomics.

### AT2 to AT1 transition

Cell-cell and cell-matrix interactions play major roles during lung development, regeneration, and repair. In the distal lung, alveolar epithelial differentiation is dependent on the reciprocal communication between epithelial and mesenchymal cells. The interaction between PDGFRa^+^ FBs and epithelial cells is essential for cellular differentiation and function during alveolarization (4, 14, 17, 18, 23). Loss of beneficial matrix-producing fibroblasts affects alveolar repair in aged and IPF lungs, suggesting the importance of fibroblast subpopulations in maintaining the normal epithelial cell niche (42). In a recent study of perinatal hyperoxia exposure we demonstrated trans differentiation of myo to lipofibroblasts which was associatedwith impaired AT2 to AT1 transition (43). In the adult lung the AXIN2 cell-lineage derived PDGFRα positive MANCs support AT1 and AT2 differentiation and function (15). *In vitro,* PDGFRa^+^ FBs support formation of alveolar organoids when cultured with primary AECs (42, 43, 45, 52). In the GATA6^PDGFRAΔ/Δ^ lungs we observed an increase in AT2 cells. Expression of Axin2 and nuclear YAP in these AT2 cells suggest that loss of GATA6 in PDGFRa fibroblasts impairs AT2 to AT1 transition (45, 46). In the present organoid studies WNT signaling restored the ability of GATA6^PDGFRAΔ/Δ^ fibroblasts to support AT1 differentiation. Recent studies demonstrate that Wnt5 a is important for AT1 differentiation and regulates AT2/AT1 balance (50). However, Wnt5a was upregulated at PN28 while the number of AT2 cells was still significantly higher, suggesting that WNT5a alone is not sufficient for AT2 to AT1 differentiation. In the GATA6^PDGFRAΔ/Δ^ lungs we observed detachment of AT1 and AT2 cells from the basement membrane likely related to the lack of ECM. While collagen was restored by PN28 the scaffolding protein CRTAP was still missing. Whether restoring the ECM will provide the missing cues for transitional AT2 cells to differentiate into AT1 cells remains to be investigated. Collectively these data demonstrate that GATA6 regulates matrix fibroblast differentiation and function. We identified a GATA6/WNT signaling pathway regulating SCMF differentiation and paracrine signaling requirements for AT2/AT1 differentiation.

## Supplemental Materials and Methods

### Animals

Mice were maintained on campus at the CCHMC animal facility. Mice were on a 12hrs:12hrs dark/light cycle and received food and water *ad libitum*. Pups were euthanized with 50 μl of triple sedative solution (67mg/ml ketamine + 3.3 mg/ml xylazine + 1.7 mg/ml acepromazine) by intraperitoneal injection and a method of secondary euthanasia was employed (exsanguination by severing the carotid artery). Adult mice used in the PN28 & PNX studies were euthanized in the same manner but received 100 ul of triple sedative solution.

### Cell culture

Human IMR90 fibroblasts (ATCC^®^ CCL-186^™^) were cultured in growth medium (DMEM Hams F12; 10% FBS; 0.1% penicillin and streptomycin; 0.1% gentamycin and amphotericin). Transfections with and without GATA6 shRNA and over expression stable lentiviral constructs (Origene) were carried out in media without antibiotics. Polybrene was used as the transfection reagent. Control cells for shRNA were transfected with scrambled shRNA.

### Morphometrics

Morphometric point intersection was carried out using a semi-automated quantification Fiji plugin (67). The alveolar simplification of airspace area on H&E stained lung sections of PN7 mice was quantified. Three to five sections from all five lobes of the lungs were analyzed. Non-parenchymal tissue (arteries, veins, and major airways) and exudates were excluded using the freehand selection tool. The Pref value was calculated by deleting points that fell within non-parenchymal tissue from the 64 point grid overlaid on the image. Psep is calculated by counting all the 1-pixel particles that fall within the septal tissue and overlaid grid excluding non-parenchymal tissue. I was counted as the number of times that the edge of the lung tissue intersected an 8 x 8 grid of 567 pixels. The unit length of the images was calculated as d = (141.75 * x)/680 um, and was used to calculate the mean linear intercept (MLI = 2 * d * (∑Pref - ∑P_sep_)/∑I). Alveolar surface area (SA Density = (2 * ∑I)/(d * ∑P_sep_)), volume (∑P_sep_/∑P_ref_), and mean transsectional wall length (2 * d * ∑P_sep_/∑I) were also calculated.

### Immunofluorescence

Immunofluorescent staining was performed on 5-µm slides sectioned from paraffin-embedded lung tissue blocks. Slides were deparaffinized in xylene, rehydrated in a series of graded ethanol and washed in 1X PBS. Antigen retrieval was performed in 10 mM citrate buffer (pH 6.0) when required. Non-specific antigens were blocked in 4% normal donkey serum in PBS with 0.1% Triton X-100 (PBST) for 2 hours. Slides were incubated in primary antibodies (Supplemental Table 1) diluted in blocking buffer overnight at 4°C. After washing in PBST, slides were incubated in fluorescent secondary antibodies (1:200) and DAPI (1µg/ml) diluted in blocking buffer for 1 hour at room temperature. Slides were subsequently washed in PBST and mounted in Prolong Gold (Thermo Fisher Scientific). Z-stack images were captured on a Nikon AR1 inverted confocal microscope and further analyzed using Nikon Elements software. Antibody staining was quantified on Nikon Elements using a previously established protocol (68). Antibodies used are listed in Supplemental Table 1.

### Cell isolation, RNA purification and quantification

PDGFRα^+^ cells were isolated from single cell suspensions by positive selection using magnetic activated cell sorting (Miltenyi Biotec, MACS), following manufacturer’s instructions. Anti-Mouse CD326 (EpCAM) isolated epithelial cells, and Anti-Mouse CD140α (PDGFRα) isolated PDGFRa^+^ FBs (Miltenyl Biotec, Auburn, CA). RNA was extracted from MACS-sorted cells or cells harvested from IMR90 cell culture using the RNeasy Mini Kit (Qiagen, No. 74104). RNA quality was verified using a spectrophotometer (Thermo Scientific Nanodrop). RNA was reverse-transcribed into single-stranded complementary DNA (cDNA) using the Verso complementary DNA synthesis kit (LifeTechnologies, AB-1453). Quantitative PCR was conducted using Taqman primers and Taqman master mix (LifeTechnologies) and measured using StepOnePlus Real Time PCR system (Thermo Fisher Scientific). Primers used are listed in Supplemental Table 2.

### Western blot

IMR90 cells treated with and/or without GATA6 shRNA and over-expression plasmids were harvested, and protein abundance was determined by western blot analysis following standard protocols. Briefly, the lysates were electrophoresed and then transferred onto a PVDF membrane. The membranes were blocked and incubated with primary antibodies overnight, which are listed with their dilutions in Supplemental Table 3. Enhanced chemiluminescent reagent was used to detect the proteins and imaged the blots using BioRad ChemiDoc Tough Imaging System. The target proteins were normalized with beta actin. Target protein expression was quantified by densitometric analysis of scanned blots using ImageJ software. Briefly, western blot bands were selected using the rectangular selection tool and defined each band by numbering them. A histogram was generated for each band and a straight-line selection tool was used to draw the line on the bottom of picks to make it close completely. Wand tool from ImageJ generated the Area and percent values for the histograms and generated the excel sheets, which was used to analyze the fold change of protein of interest compared to beta actin. The percent values of protein of interests were normalized over percent values of beta actin. Fold change was calculated over control samples.

### Collagen Contraction Assay

Collagen pellets were generated following previously established protocols (69-71). Collagen pellets were prepared in a 12 well tissue culture plate that was coated with 1% BSA in media and incubated for 2 hrs at 37°C. After removal of media, 1ml of 1 mg/ml type I collagen (BD Biosciences) was added and allowed to polymerize for 16 hrs at 37°. The resulting collagen matrix was detached from the sides of the well using a sterile 10μl pipette tip. IMR90 cells transfected with and/or without shRNA and over expression plasmids were seeded on the free-floating collagen matrix at 100,000 cells in 10% FBS/DMEM Hams F12 media. Pellets were cultured for 24 hrs and then imaged using a Leica DFC7000 T camera attached to a Leica MZ16 FA fluorescent stereomicroscope. Contraction was measured with Image J software by calculating the circumference of the pellet and dividing by the circumference of the well, calculating percentage contraction. Multiple t-test analysis was performed using GraphPad Prism software.

### Chromatin Immunoprecipitation

Chromatin Immunoprecipitation (ChIP) was performed using low cell ChIP kit (abcam), according to the manufacture’s protocol. IMR 90 cells stably transfected with GATA6 shRNA and overexpression plasmids were immunoprecipitated over night at 4^0^C using EZH2 antibody (Cell Signaling Technology). Immune complexes were pulled down using IgG coated wells. DNA cross-links of the immune complexes was reverted and DNA was isolated. The retrieved DNA was analyzed by qPCR using TCF21 primers (Supplemental Table 4) and fold enrichment was calculated. Normal human IgG served as negative control.

### Transmission Electron Microscopy

Postnatal day 7 and day 28 mouse lungs were inflation fixed with 2% paraformaldehyde (Electron Microscopy Sciences (EMS), Hatfield, PA), 2% glutaraldehyde (EMS), 0.1% calcium chloride (Fisher Chemcials, Fairlawn, NJ) in 0.1 M sodium cacodylate (EMS) buffer (SCB), pH 7.2, under 25 cm water pressure, followed by post fixation with fresh fixative at 4 °C for up to 2 days. Lung lobes were cut into 1–2 mm blocks and processed for transmission electron microscopy as previously described (72, 73). Ultrathin sections were cut using a Reichert-Jung Ultracut E ultramicrotome (Reichert-Jung, Austria), and EM images were digitally acquired using a Hitachi H-7650 transmission electron microscope (Hitachi High Technologies USA, Schaumburg, IL) equipped with a CCD camera (Advanced Microscopy Techniques. Woburn, MA) at 80 kV.

### RNA-seq and bioinformatic analysis

MACS-sorted PDGFRA^+^ (CD140^+^) fibroblasts and EPCAM^+^ epithelial cells from either control or GATA6^PDGFRA,1/,1^ lungs at PN7 were prepared for bulk RNA-seq. RNA sequencing was performed by Genewiz from Azenta Life Sciences. RNA-seq FASTQ files were aligned using Bowtie to mouse genome version mm10 (74). Raw gene counts were obtained using Bioconductor’s Genomic Alignment, which were subsequently made into normalized FPKM values using Cufflinks (75, 76). DeSeq (Bioconductor) was used to calculate differential gene expression from raw expression values. Genes were deemed differentially expressed if they satisfied the following requirements: gene has a fold change >2, binomTest p-value <.01, and RPKMs >1 in 2 of the 3 replicates in at least one condition being compared. Gene patterns were determined by comparing differential expressed genes from all three time points. Genes that were significantly changed or unchanged at the same time points were grouped together. Heatmaps of genes in particular patterns were z-score normalized and generated using Partek Genomics Suite (http://www.partek.com/pgs). Gene enrichment analysis was carried out using ToppGene’s ToppFun, and functional enrichments within each profile were identified and all profiles were compared to each other using Toppcluster (77). Pvalues of functional enrichment hits where –log10 transformed for graphical visualization. Fold changes of any WNT signaling pathway genes were visualized in a heatmap generated by pheatmap (https://cran.r-project.org/web/packages/pheatmap/index.html).

### Statistics

An Unpaired Student’s two-tailed t-test was used to determine the significance between two groups. A One-way ANOVA followed was used to determine significance between three groups. GraphPad Prism 9.0.0 was used to calculate statistical differences and for creation of the associated graphs. P<0.05 was considered statistically significant.

### Study approval

Mice are housed in a pathogen-free, an Association for Assessment and Accreditation of Laboratory Animal Care (AAALAC) accredited facility at Cincinnati Children’s Hospital Medical Center (CCHMC). All animal protocols used in this study were approved by the Institutional Animal Care and Use Committee (IACUC) at CCHMC.

## Author Contributions

MGU: designed, conducted, and analyzed in vitro experiments with IMR90 cells, contributed to lung harvests for RNA; contributed immunohistochemistry staining and analyzing morphometric analysis; contributed to figures and writing the manuscript.

JG: conducted animal breedings, genotyping, partial pneumonectomy surgeries, and animal harvests; designed, conducted, and analyzed all organoid studies, contributed to morphometric analysis, conducted most of the immunohistochemistry and confocal imaging, generated figures, contributed to writing the manuscript

MR: conducted gene expression studies in primary cells; designed, conducted, and analyzed the collagen pellet experiments, contributed to immunohistochemistry, contributed to figures, and writing the manuscript.

CN: conducted and analyzed all electron microscopy experiments.

DM: contributed to immunohistochemistry and morphometric experiments for PN7.

MG: performed functional enrichment analysis on the RNA-seq data and generated the associated PC and volcano plots and extracted WNT signaling pathway genes to create the heatmap.

AKP: designed, funded, and supervised all research studies, analyzed data, generated the graphical abstract, generated all figures and wrote the manuscript.

## Acknowledgement

This work was supported by NIH grants R01 HL131661 (to MGU, JG, MR and AKTP), T32 HL007752 (to MR), U01 HL122642 (MG, AKTP) LungMAP1, U01 HL122638 (MG, AKTP) LungMAP2, U01 HL134745 (MG, AKTP) PCTC. Additional internal funds were from the Translational Fibrosis Academic and Research Committee funding CCHMC, and the BPD Academic and Research Committee funding CCHMC. We would like to thank Dr. Roy Morello for his generous gift of the CRTAP antibody.

---

**Supplemental Table 1.** List of primary antibodies used in this study

| **Immunofluorescent Antibody** | **Source** | **Catalog #** | **Dilution** | **Antigen Retrieval** |
| --- | --- | --- | --- | --- |
| aSMA | Abcam | A5228 | 1:10,000 | Citrate |
| ADRP | Abcam | Ab52356 | 1:200 | Citrate |
| PDGFRA | R+D Systems | AF1062 | 1:50 | Citrate |
| FN1 | Abcam | Ab23750 | 1:100 | Citrate |
| CRTAP | Gift from<br>Dr. Roy Morello | N/A | 1:200 | Citrate |
| ECAD | R+D Systems | MAB7481 | 1:2000 | None |
| AGER | R+D Systems | AF1145 | 1:200 | Citrate |
| HOPX | Santa Cruz | Sc-30216 | 1:100 | Citrate |
| SPC | Seven Hills | R458 | 1:100 | Citrate |
| YAP | Santa Cruz | Sc-101199 | 1:50 | Citrate |

**Supplemental Table 2:** List of primary antibodies used for western blot

| Antibody | Source | Catalog # | Dilution |
| --- | --- | --- | --- |
| GATA6 | R&D Systems | AF1700 | 1:500 |
| EZH2 | Cell signaling Technology | D2C9 | 1:500 |
| Beta actin | Seven Hills | LMAB-C4 | 1:1000 |

**Supplemental Table 3:** List of Taqman primers used for qPCR

| Gene | TaqMan probe catalog # |
| --- | --- |
| *Pdgfra* | Mm00440701 |
| *Gata6* | Mm01235633 |
| *Tcf21* | Mm00448961 |
| *Plin2* | Mm00475794 |
| *Pparg* | Mm00440940 |
| *Ctsk* | Mm00484039 |
| *Wnt5a* | Mm00437347 |
| *Axin2* | Mm00443610 |
| TCF21 | Hs00162646 |
| PLIN2 | Hs00605340 |
| WNT2 | Hs00608224 |
| WNT5A | Hs00998537 |
| GATA6 | Hs00232018 |
| ELN | Hs00355783 |
| COL1A1 | Hs00164004 |
| ACTA2 | Hs00426835 |
| FN1 | Hs01549976 |

**Supplemental Table 4:**
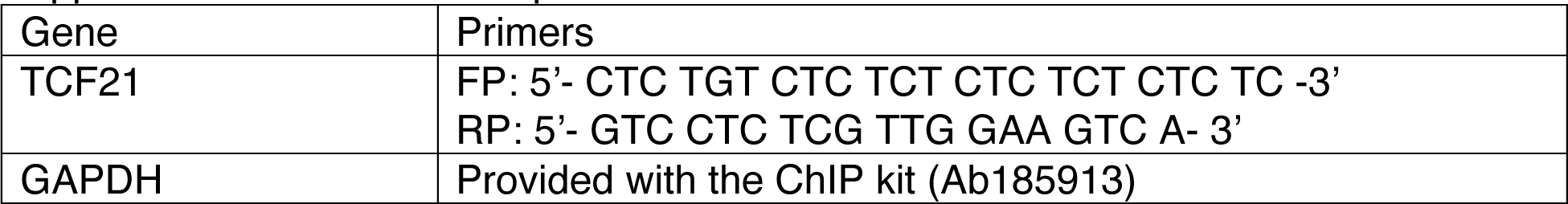
List of primers used for ChIP

**Supplemental figure S1:**
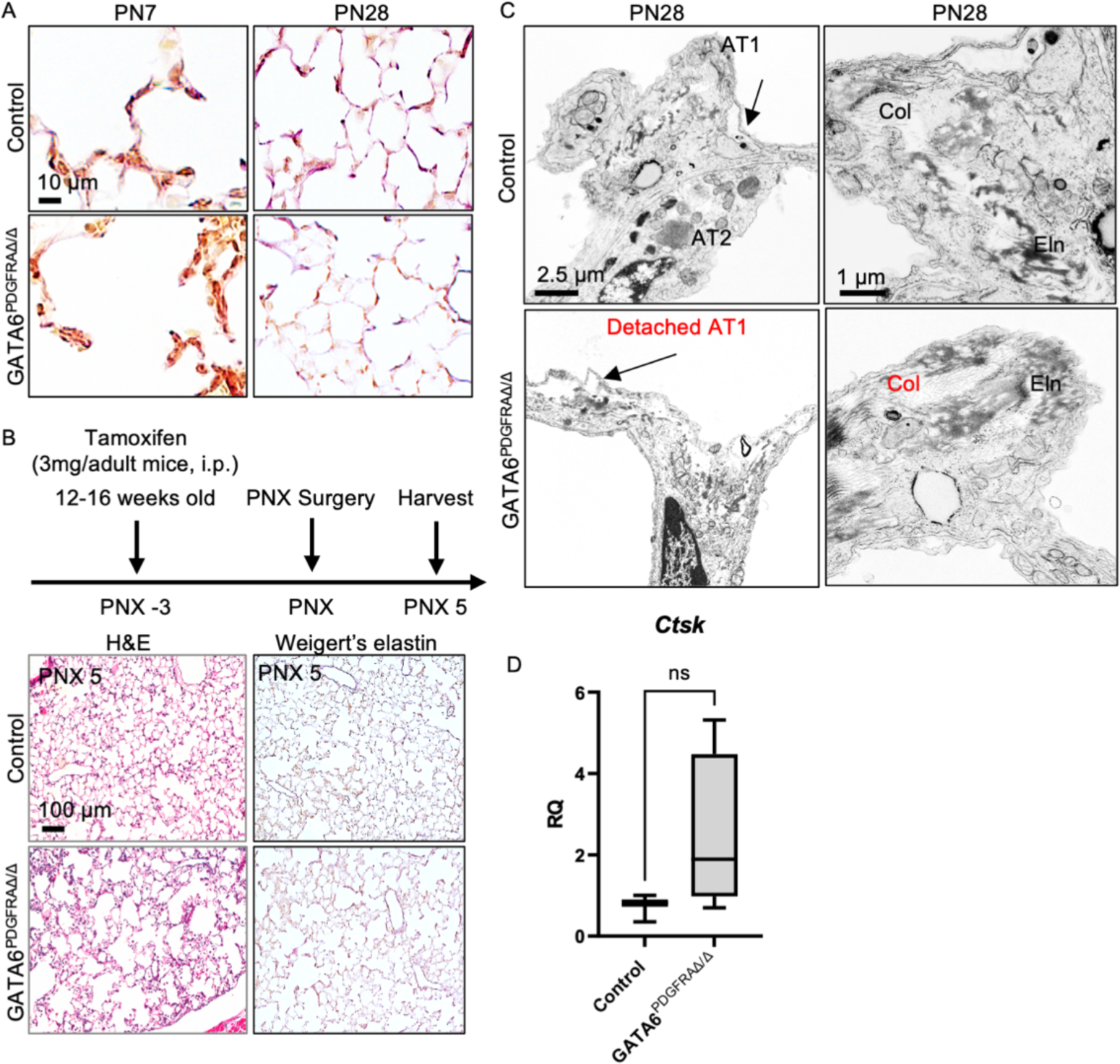
**(A)** Weigert’s elastin staining showed less elastin and collagen deposition in GATA6^PDGFRAΔ/Δ^ lungs at PN7 septal tips, while no change at PN28. Scale bars= 10µm. **(B)** Schematic representation of tamoxifen administration and lungs harvest timings post-PNX, H&E and Weigert’s elastin D5 post-injury showed alveolar simplification as observed during developmental alveolarization. Scale bars= 100 µm. **(C)** Transmission electron microscopy of GATA6^PDGFRAΔ/Δ^ lungs showed restoration of collagen fibers and reattachment of AT1 cells to the basement membrane at PN28. Scale bars= 2.5 µm, 1 µm. (D) A trending increase in Cathepsin K gene expression was observed in GATA6^PDGFRAΔ/Δ^ fibroblasts at PN7, likely causing the degradation of collagen at PN7. n=3-4, Two-tailed Student’s t-test, ns-not significant.

**Supplemental figure S2:**
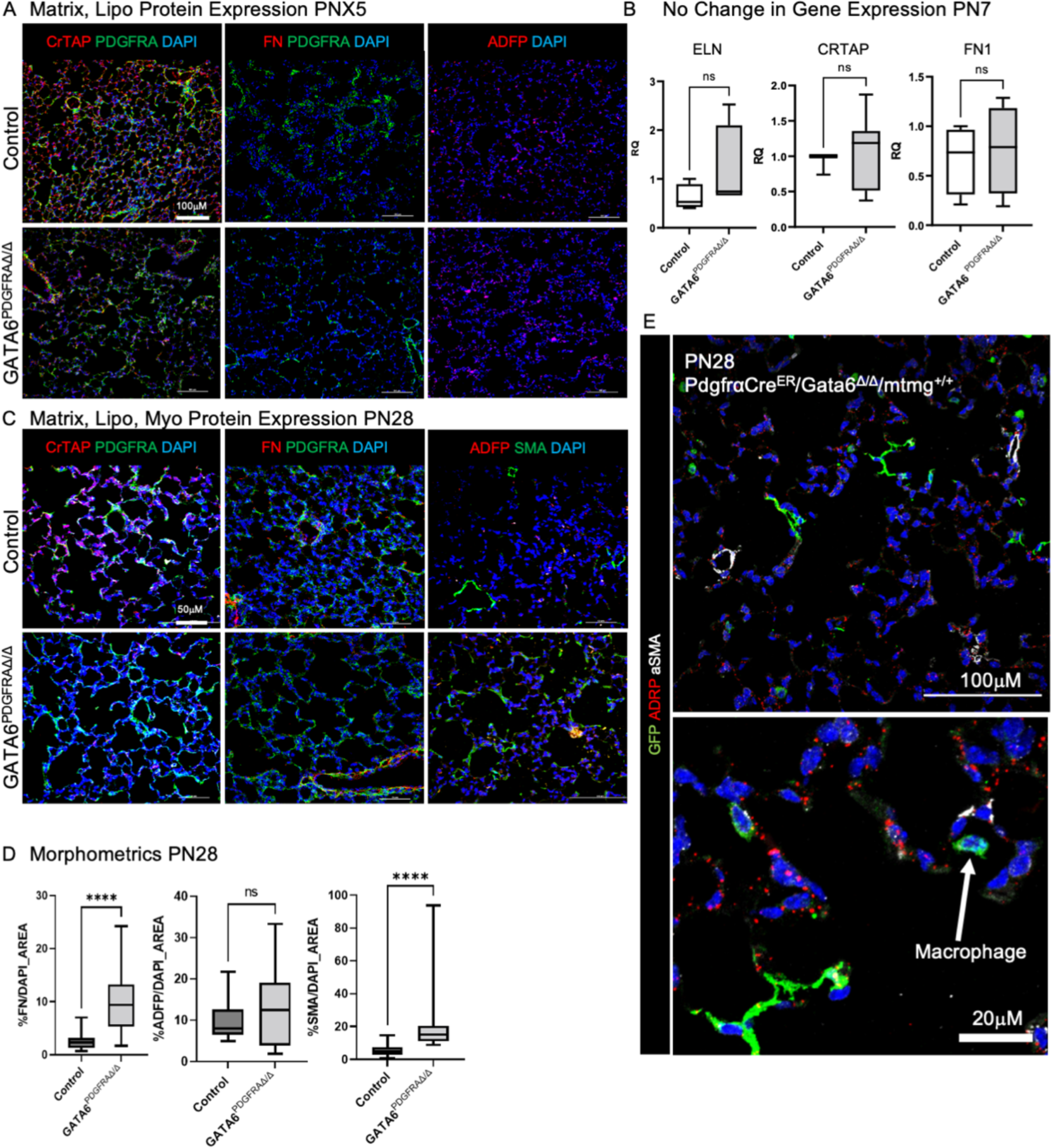
**(A)** Immunostaining showed reduced matrix protein expression of CrTAP and FN, increased ADRP at D5 post-PNX in GATA6^PDGFRAΔ/Δ^ lungs as seen in developmental alveolarization. Scale bars=100µm. **(B)** MACS-sorted PDGFRa+ cells showed no change in matrix fibroblast-specific gene expressions (Crtap, Eln & Fn) at PN7. n=3-6, Two-tailed Student’s t-test was used, ns-not significant. **(C)** Immunostaining of CrTAP, FN & ADFP in control and GATA6^PDGFRAΔ/Δ^ lungs at PN28. **(D)** Quantification was done using Nikon Elements. ADFP was unchanged, FN and aSMA were increased at PN28. n=3-4. Two-tailed Student’s t-test was used, ****P<0.0001, ns-not significant. **(E)** PN28 immunofluorescence for PDGFRa-GFP lineage trace of SCMF and colocalization of aSMA (white) and ADRP (red) in GATA6^PDGFRAΔ/Δ^ lungs. At PN28 most GFP is found in macrophages as they clear the SCMF. Remaining lineage traced fibroblasts tare neither aSMA nor ADRP positive.

**Supplemental figure S3:**
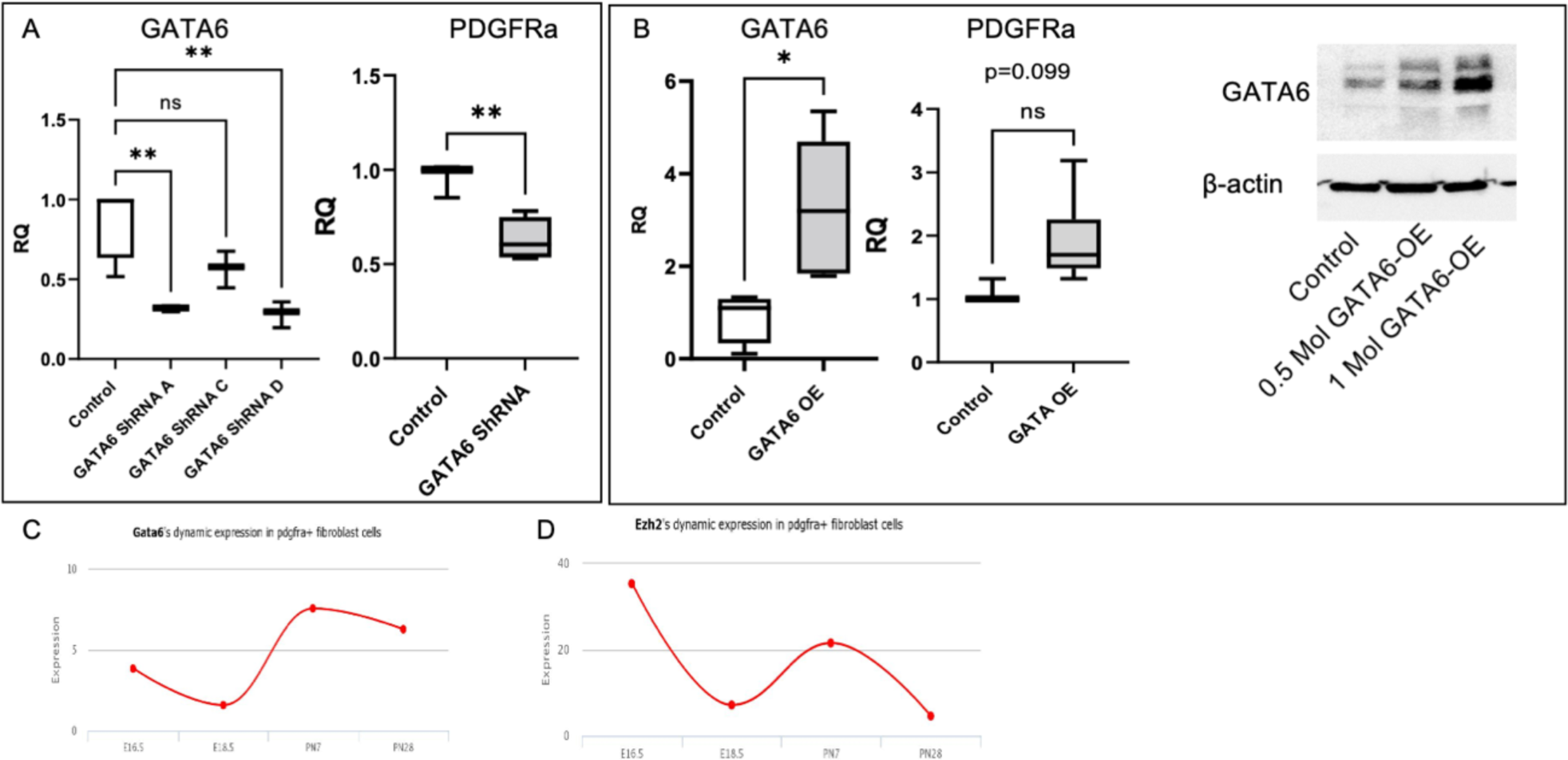
**(A)** IMR90s were transfected with lentiviral particles containing GATA6 ShRNA. Transfection efficiency was confirmed by qPCR. n=3. Ordinary one-way ANOVA was done. **p<0.005, ns- not significant. PDGFRA gene expression was downregulated with GATA6 ShRNA, N=3-4, Two-tailed Student’s t-test was used. **p=0.0100 **(B)** GATA6 over expression efficiency standardization of the multiplicity of infection (MOI) of the lentiviral overexpression plasmid was confirmed using qPCR and Western blotting. Two different MOIs were carried out and 1MOI significantly increased in GATA6 protein expression and used that for all the further experiments. Beta actin was used as the loading control. PGDFRA gene expression was trending up with GATA6 OE, n=4-6. Two-tailed Student’s t test was used. *p<0.05, ns-not significant. **(D)** LGEA graph showing dynamic expression changes in GATA6 gene expression during alveolarization **(E)** Similar pattern was identified by LGEA for EZH2 expression during alveolarization.

**Supplemental figure S4:**
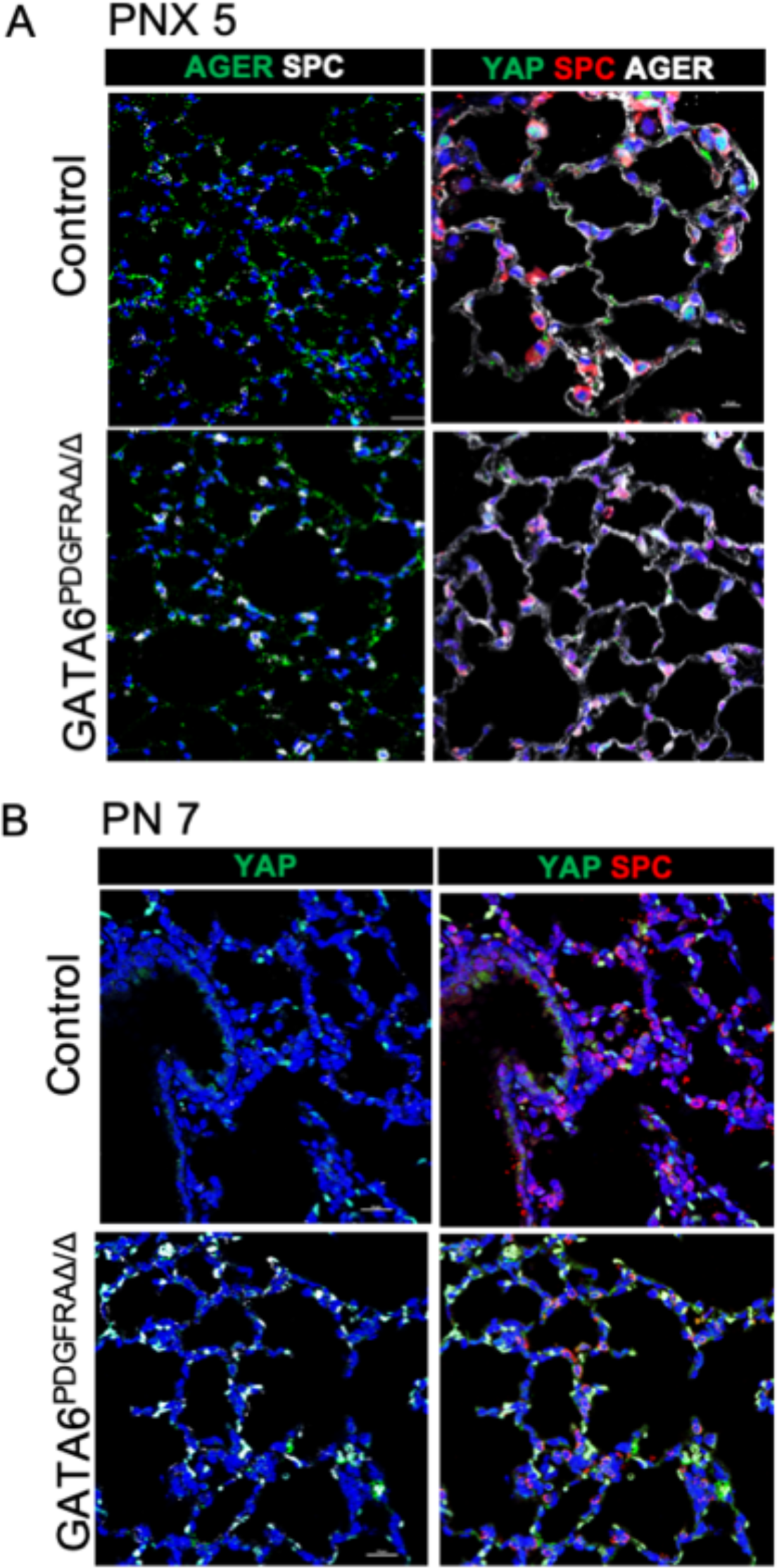
**(A)** Immunofluorescence showed increase in SPC^+^AT2 cells and decrease in AGER+ AT1 cells 5 days post PNX. Immunofluorescence of YAP was also performed at D5 post PNX. **(B)** Immunofluorescence of YAP and SPC at PN7.

**Supplemental figure S5:**
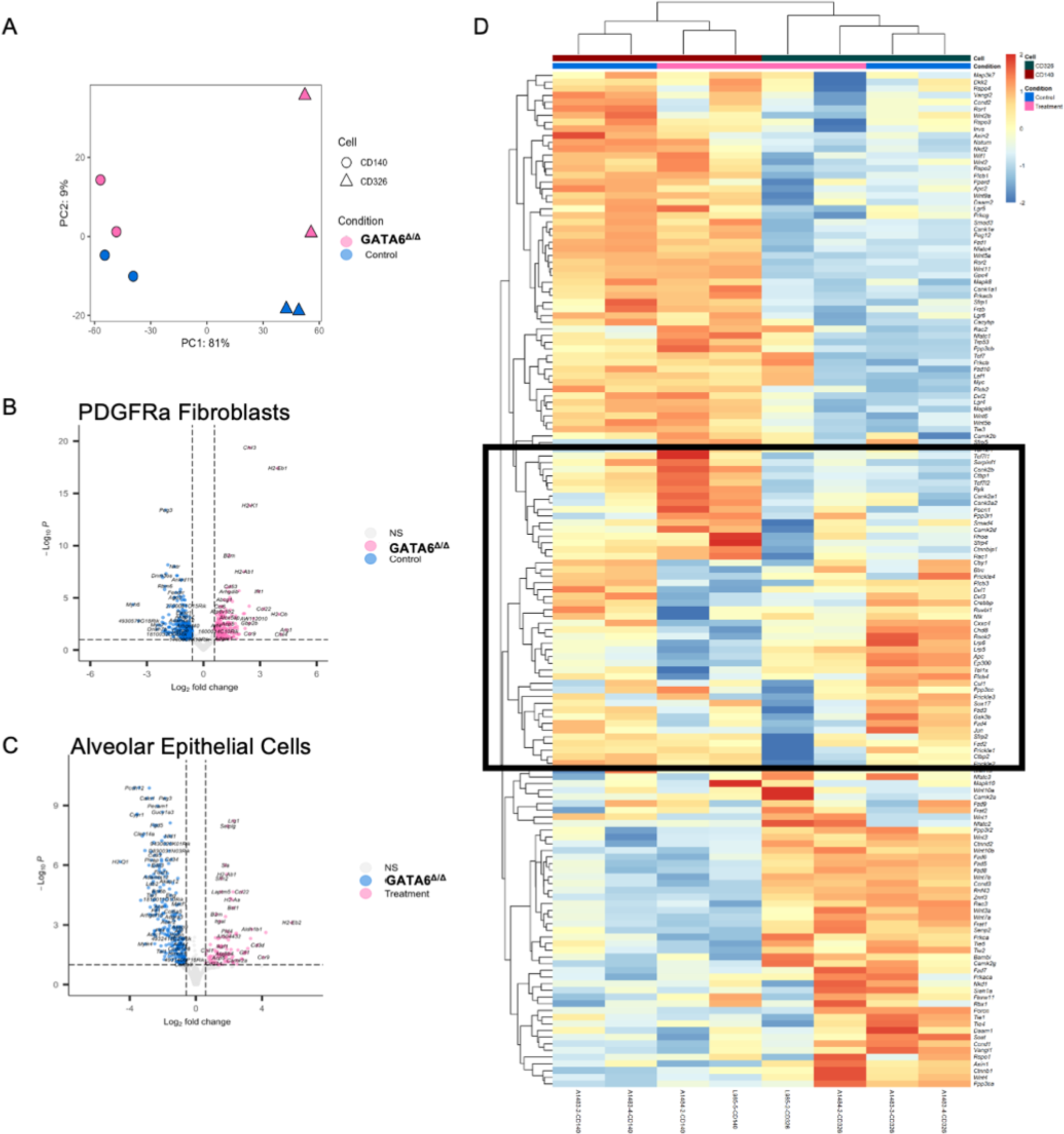
Bulk sequencing of sorted CD140+ fibroblasts and Epcam+ epithelial cells of control and GATA6^PDGFRAΔ/Δ^ lungs. **(A)** Principal component analysis of PDGFRa+ fibroblasts (CD140a+) and epithelial cells (CD326+) control and GATA6^PDGFRAΔ/Δ^ lungs. **(B)** Volcano plots **(C)** Heatmap with 152 WNT related genes from the KEGG pathway.

